# In vitro dimerization of human RIO2 kinase

**DOI:** 10.1101/553800

**Authors:** Frédérique Maurice, Natacha Pérébaskine, Stéphane Thore, Sébastien Fribourg

## Abstract

RIO proteins form a conserved family of atypical protein kinases. RIO2 is a serine/threonine protein kinase/ATPase involved in pre-40S ribosomal maturation. Current crystal structures of archaeal and fungal Rio2 proteins report a monomeric form of the protein. Here, we describe three atomic structures of the human RIO2 kinase showing that it forms a homodimer *in vitro*. Upon self-association, each protomer ATP-binding pocket is partially remodelled and found in an apo state. The homodimerization is mediated by key residues previously shown to be responsible for ATP binding and catalysis. This unusual *in vitro* protein kinase dimer reveals an intricate mechanism where identical residues are involved in substrate binding and oligomeric state formation. We speculate that such an oligomeric state might be formed also *in vivo* and might function in maintaining the protein in an inactive state and could be employed during import.

## Introduction

Ribosome biogenesis is a complex mechanism initiating in the nucleolus with the transcription of a long ribosomal RNA precursor (pre-rRNA) by RNA polymerase I, and ending in the cytoplasm after interconnected maturation events that include methylation, nucleotide pseudouridylation, endo- and exonucleolytic cleavage and a mandatory transit through the nuclear pore ^1,2^. Some of the 200 associated factors (AF) involved in ribosome biogenesis ^3–8^ display characteristic sequence motifs for enzymes such as Ser/Thr kinases, GTPases, RNA helicases, and are probably involved in the many checkpoint controls. The three proteins forming the RIO kinases family, namely Rio1, Rio2 and Rio3, have a distinct role in cytoplasmic maturation of the pre-40S pre-rRNA ^9–13^. These proteins form a family of Ser/Thr kinases built around a central protein kinase domain but do not exhibit further sequence similarity with other eukaryotic protein kinases (ePKs), and as such have been classified as atypical protein kinases ^14,15^. In yeast, *RIO1* and *RIO2* are essential genes and their products, Rio1 and Rio2, associate to ribosomal pre-40S particles ^11,13,16^ and are involved in D-site cleavage *in vivo* ^11,13,16,17^. Rio1 is additionally involved in cell cycle progression and rDNA stability ^18–20^. The individual depletion of Rio1 and Rio2 leads to an accumulation of the 20S pre-rRNA in yeast ^16,17,21,22^ and 18S-E pre-rRNA in human cells ^9,23^. In humans, hRIO3 is exclusively cytoplasmic and forms part of the pre-40S particles with association to the 18S-E pre-rRNA. hRIO3 deletion leads to 21S rRNA accumulation in humans ^10^.

The presence of the human RIO2 protein (hereafter hRIO2), but not its activity, is necessary for the recycling of the hENP1 factor from the cytoplasm to the nucleus. In contrast, hRIO2 activity is required for the release of hDIM2, hLTV1, hNOB1 and hRRP12, from the 18S-E pre-rRNA ^13,24^. In addition, hRIO2 phosphorylates hDIM1 and hRIO2 auto-phosphorylates *in vitro* ^13,25^. In yeast, Rio2 catalytic activity is not required for its association to the pre-40S particle but rather for its release ^26,27^. Altogether, these observations led to the proposal that Rio2 is rather an ATPase than a protein kinase. Finally, the position of Rio2 on the pre-40S particle has been established at the P-site close to domain IV of Tsr1, where both proteins block the A- and P-sites on the 40S subunit ^28,29^.

The structure of the archaeal *Archeoglobus fulgidus* Rio2 (afRio2) and *Chaetomium thermophilum* Rio2 (ctRio2) have been solved in the apo and ATP-bound states ^14,26,30^. These atomic models reveal the domain organization of Rio2 proteins and give insights into the enzymatic reaction mechanism. Each of these structures, however, describes Rio2 as a monomer. Here, we report that hRIO2 can form a homodimer in solution. We solved the crystal structure of the homodimer of hRIO2 and observe that several invariant or highly conserved residues are involved in the dimer interface. In the hRIO2 homodimer, the ATP-binding site is modified in a conformation preventing ATP binding. Moreover, residues involved in the homodimer interaction are also required for ATP-binding and catalysis. From the structure, the enzymatic activity and the dimer formation would be mutually exclusive, and dimerization is incompatible with pre-40S particle association. In absence of *in vivo* evidence so far, we propose that this dimeric state could be transient and could represent a conformation used for protein import to the nucleus.

## RESULTS

### Solution analysis of human RIO2

Rio2 is a phylogenetically conserved protein whose size varies from approximately 30 kDa in the archaea *A. fulgidus* to about 60 kDa in humans. The proteins are organized around a N-terminal winged helix domain followed with a classical kinase domain architecture with a N- and C-terminal lobes. In higher eukaryotes, the C-terminus contains additional elements such as a Nuclear Export Signal (NES) ^31,32^. The full-length hRIO2 protein was found to be poorly expressed and highly unstable, and therefore various C-terminally truncated constructs were first engineered in order to remove natively unfolded regions of the protein. We also generated constructs that removed the unfolded loop located between residues 131 to 146, since this loop promoted instability of the protein and strong nonspecific binding to Ni^+^-NTA and Co^2+-^NTA resin. The enzymatic activity of the truncated samples was controlled using a phosphorylation assay as described in Zemp et al. 2009 ^13^, and compared to full-length hRIO2 and ctRio2 (Figure S1). A phosphorylation activity was detected for all constructs demonstrating that the truncations are not impaired in the enzymatic activity.

To evaluate the oligomeric state of hRIO2 in solution, we used sedimentation velocity analytical ultracentrifugation (SV-AUC). To this end, we purified to homogeneity an hRIO2 construct encompassing residues 1-321, with a deletion of the unfolded loop (residues 131-146) ^26,30^. As a control, we used the full-length ctRio2 ^26^. Protein concentrations varied from 3.75 μM to 30 μM for ctRio2 and from 3.37 μm to 27 μM for hRIO2 in SV-AUC (Fig. 1A to 1D). For ctRio2, the continuous c(s) distribution plots showed one major sedimentation species with a sedimentation coefficient of 3.16 S and an estimated molecular mass of 44.80 kDa, which is close to the theoretical value of 47.24 kDa for the monomer. These results indicate that ctRio2 behaves as a single monomeric species at all tested protein concentrations (Fig. 1A and 1C and Table 1). For hRIO2(1-321)*Δ*(131-146), the c(s) analysis revealed a concentration-dependent behavior, with a large majority (> 94 %) of monomer at lowest concentrations (3.375 μM) and an increasing proportion of dimer at higher concentrations (> 20 % at 27 μM) (Fig. 1B and 1D and Table 1). In contrast to the reported atomic models of afRio2 and ctRio2, these sedimentation velocity experiments reveal the ability of hRIO2(1-321)*Δ*(131-146) to form homodimers.

**Fig. 1:**
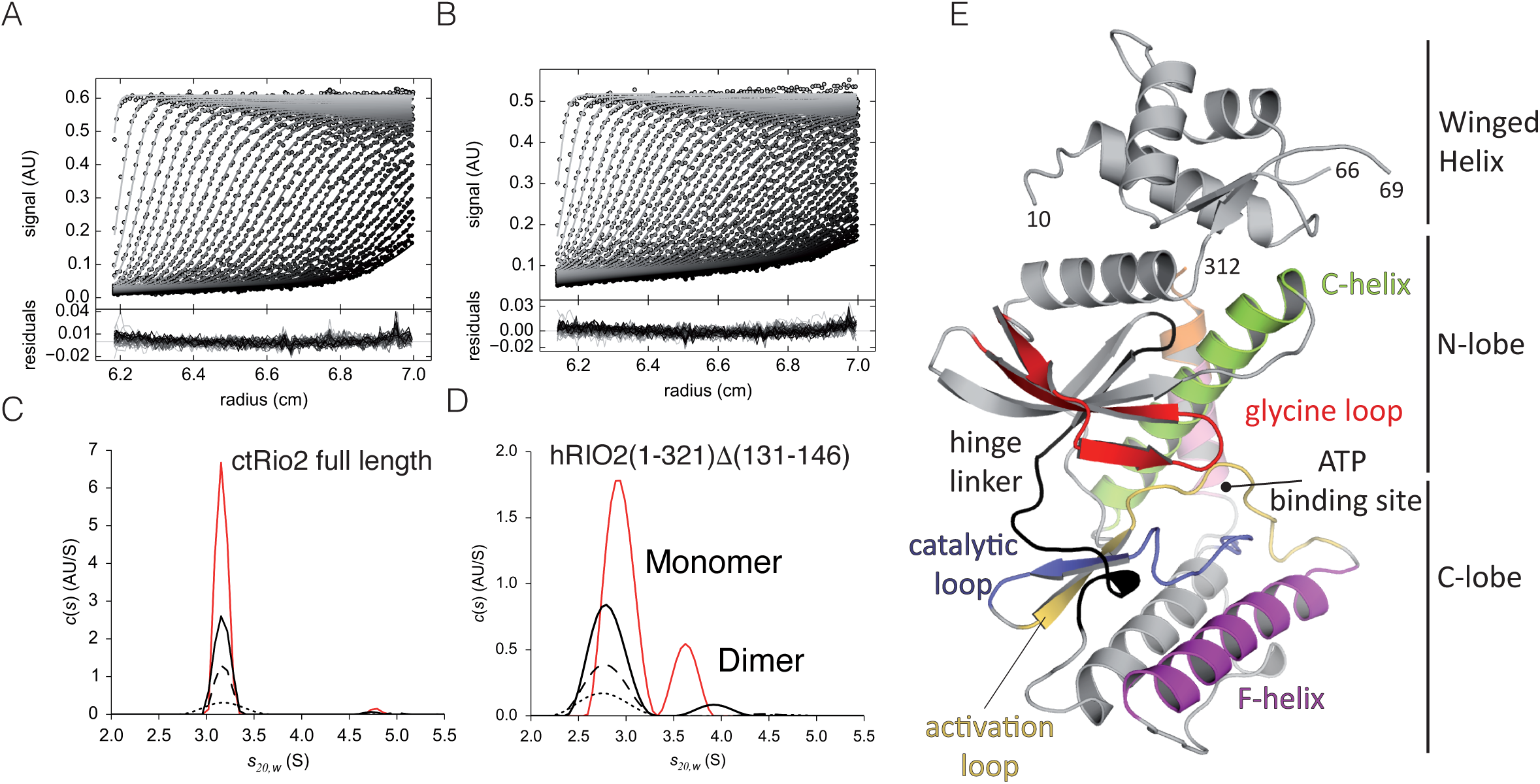
Analytical UltraCentrifugation (AUC) studies and structure of human RIO2 kinase. AUC experiments were performed at 4 °C with 50,000 rpm. For clarity, only one of three acquisitions is shown, corresponding to about 15 min time intervals. Continuous size (c(s)) distribution (y-axis) are plotted as a function of sedimentation coefficient (x-axis) in Svedberg (S). Sedimentation coefficient distributions *c*(s) were determined from Sedfit/Sedphat ^50,52^ analysis of absorbance data at 280 nm. **A)** Superposition of experimental and fitted sedimentation velocity profiles (top), and their differences (bottom) for ctRio2 at 15 µM and **B)** hRIO2(1-321)*Δ*(131-146) at 13.5 µM. **C)** ctRio2 full length at 30 µM (orange line), 15 µM (black line), 7.5 µM (dashed line) and 3.75 µM (dotted line) appeared at all concentrations as a monomer. Sedimentation velocity data were plotted using the software GUSSI. **D)** hRIO2(1-321)*Δ*(131-146) samples at 27 µM (orange line), 13.5 µM (black line), 6.75 µM (dashed line) and 3.375 µM (dotted line) were analyzed. hRIO2(1-321)*Δ*(131-146) appears as a dimeric upon increasing concentrations. **E)** Structure of the hRIO2(1-353)*Δ*(131-146) encompassing residues 1 to 353. The winged helix is located at the N-terminus, the two kinases lobes are labelled and connected by a hinge linker loop (in black). The so-called C-helix (light green), F-helix (magenta), glycine loop (red), catalytic loop (blue) and activation loop (yellow) are also shown and labelled accordingly to the canonical representation ^15^. C-terminal residues to the F-Helix are coloured in pink and orange. This and all subsequent structural Figures were generated with PyMOL ^58^.

**Table 1.** Analytical ultracentrifugation studies of hRIO2(1-321)Δ(131-146) and full length ctRio2.

| **[hRIO2]** | **% monomer** | **s<sub>20,w</sub> monomer** | **% dimer** | **s<sub>20,w</sub> dimer** | **% oligomer** |
| --- | --- | --- | --- | --- | --- |
| 3.37 μM | 94.7 | 2.76 S | 0.7 | 4.28 S | 4.6 |
| 6.75 μM | 95.7 | 2.78 S | 1.7 | 4.35 S | 2.6 |
| 13.5 μM | 91.2 | 2.80 S | 8.5 | 3.92 S | 0.3 |
| 27 μM | 79.8 | 2.93 S | 20.2 | 3.62 S | 0 |
| **[ctRio2]** | **% monomer** | **s<sub>20,w</sub> monomer** | **% dimer** | **s<sub>20,w</sub> dimer** | **% oligomer** |
| 3.75 μM | 96.6 | 3.18 S | 0 | - | 3.4 |
| 7.5 μM | 97.3 | 3.17 S | 2.0 | 4.98 | 0.7 |
| 15 μM | 97.6 | 3.16 S | 2.0 | 4.73 | 0.4 |
| 30 μM | 96.6 | 3.16 S | 1.9 | 4.78 | 1.5 |

### Structure determination

In order to characterize this unexpected dimeric state of hRIO2, we decided to use protein crystallization as it is usually performed at high concentrations favourable for hRIO2 dimerization. For the construct encompassing residues 1-321, native diffraction datasets were collected up to 3.2 Å resolution although with crystallization reproducibility issues, limited diffraction and the failure to solve the structure by molecular replacement. In contrast, the construct with the deleted loop (residues 131 to 146), provided native and Se-Met derived protein crystals for which diffraction data were collected (Table 2). The structure of hRIO2(1-321)*Δ*(131-146) was determined by SAD phasing on Se-derived protein crystals. The refined structure was used as a model to subsequently solve the crystal structure of hRIO2(1-321) and of a longer construct, hRIO2(1-353)*Δ*(131-146). All refinement statistic for the three atomic structures can be found in the Table 2.

**Table 2.** Crystallographic data and refinement statistics.

| | hRIO2(1-321)$\Delta$(131-146) Se-Met | hRIO2 (1-321)$\Delta$(131-146) | hRIO2 (1-321) | hRIO2 (1-353)$\Delta$(131-146) |
| --- | --- | --- | --- | --- |
| Beamline | PX1 | PX1 | PX1 | PX1 |
| Space group | $P2_1$ | $P2_1$ | $P2_12_12_1$ | $P6_5$ |
| Unit cell parameters |  |  |  |  |
| a, b, c (Å) | 67.15 99.08 117.48 | 66.92 98.327 117.069 | 58.59 86.40 121.54 | 67.55 67.55 312.19 |
| $\alpha, \beta, \gamma$ (°) | 90.00 95.05 90.00 | 90.00 95.22 90.00 | 90.00 90.00 90.00 | 90 90 120 |
| Wavelength (Å) | 0.9797 | 0.9801 | 0.9788 | 0.9789 |
| Resolution range (Å) | 46.10 - 2.37 (2.43-2.37) | 41.67 - 2.1 (2.17- 2.10) | 70.4- 2.9 (3.07- 2.90) | 42.69 - 2.6 (2.69 - 2.6) |
| Total reflections | 232,087 (24,051) | 331,294 (31,840) | 47,340 (4,662) | 279,056 (31,169) |
| Unique reflections | 61,746 (6,173) | 87,771 (8,466) | 13,696 (1,224) | 24,610 (2,417) |
| R<sub>p</sub>im (%) | 4.09 (35.8) | 2.16 (33.59) | 4.06 (35.45) | 5.78 (73.21) |
| I/σI | 12.24 (2.26) | 20.59 (2.32) | 13.81 (2.62) | 8.73 (1.49) |
| Completeness (%) | 99.40 (99.37) | 98.49 (96.68) | 92.17 (88.49) | 99.98 (100.00) |
| Redundancy | 3.8 (3.9) | 3.8 (3.7) | 1.93 (1.90) | 11.3 (12.9) |
| Se sites | 40 | - | - | - |
| Wilson Plot B-factor (Å<sup>2</sup>) | 46.89 | 42.09 | 72.28 | 67.2 |
| CC 1/2 | 0.997 (0.845) | 0.999 (0.879) | 0.997 (0.744) | 0.991 (0.493) |
| FOM acentric (before /after solvent flattening) | 0.394 / 0.715 | - | - | - |
| Refinement statistics |  |  |  |  |
| R<sub>work</sub> (%) | 19.70 | 20.05 | 22.58 | 21.44 |
| R<sub>free</sub> (%) | 22.89 | 23.52 | 26.67 | 25.00 |
| No. of non-H atoms | 9,713 | 9,792 | 3,924 | 4,636 |
| Protein | 1,144 | 1,133 | 506 | 565 |
| Water | 413 | 596 | - | 61 |
| Average B factor (Å<sup>2</sup>) | 60.13 | 56.77 | 77.97 | 81.0 |
| Ramachandran (%) |  |  |  |  |
| Preferred/allowed/outliers (%) | 97.05 / 1.79/ 1.16 | 99.46 / 0.54/ 0.0 | 93.99/ 3.86 / 2.15 | 92.38 / 5.81 / 1.81 |
| Rmsd bond/angle (°) | 0.022 / 1.87 | 0.020 / 1.76 | 0.008/ 1.03 | 0.015/ 1.87 |
| Clash score | 7.25 | 5.70 | 5.28 | 4.95 |
| PDB | 6FDV | 6FDM | 6FDN | 6FDO |

Numbers under brackets refer to the highest resolution shell

### Overall structure description

The hRIO2 kinase adopts a similar fold as afRio2 (1TQP) from the archea *A. fulgidus* and ctRio2 (4GYI) from a thermophilic fungi ^26,30^. The N-terminus (residues 10 to 75) folds as a winged helix domain (Fig. 1E), which is situated adjacent to the two lobes of the Rio kinase domain (respectively residues 76-190 and 196-291). The hRIO2 structure contains the typical motifs of canonical Ser/Thr protein kinases with a C-Helix (light green), the F-Helix (magenta), the catalytic loop (in dark blue) and the glycine loop (red, also known as P-loop). The hinge linker (residues 191-195, black) involved in the recognition of the ATP adenine moiety in afRio2, connects the two lobes of the kinase domain (Fig. 1E). No electron density for ATP or any ligand is seen in the electron density map at or near the active site.

hRIO2(1-321) and hRIO2(1-353)*Δ*(131-146) have 1.2 Å root-mean-square deviation (r.m.s.d.) over 249 C-*α* carbon and differ mainly by the relative position of the C-lobe (Fig. S2A). Despite sequence identity and structure similarity with hRIO2, both ctRio2 and afRio2 display slightly different domain orientations as shown by relative r.m.s.d. of 2.5 Å (221 C-*α* carbons) and 1.5 Å (244 C-*α* carbons), respectively (Fig. S2B and S2C). Specifically, the N-terminal winged helix domains and C-terminal lobes of the kinase domain are in slightly altered orientations (Fig. S2).

### Structure of human RIO2 homodimer

In the asymmetric unit, we observe a homodimer formed by the association of two hRIO2 protomers in a head to head orientation independent of the three space groups or of the construct size we used (Fig. 2A). The F- and the C-helices form the major part of the homodimer interface without significant impact on the N-terminal winged helix domains. The F-helix of one protomer interacts with the F-helix of the other protomer in an antiparallel mode (Fig. 2B).

**Fig. 2:**
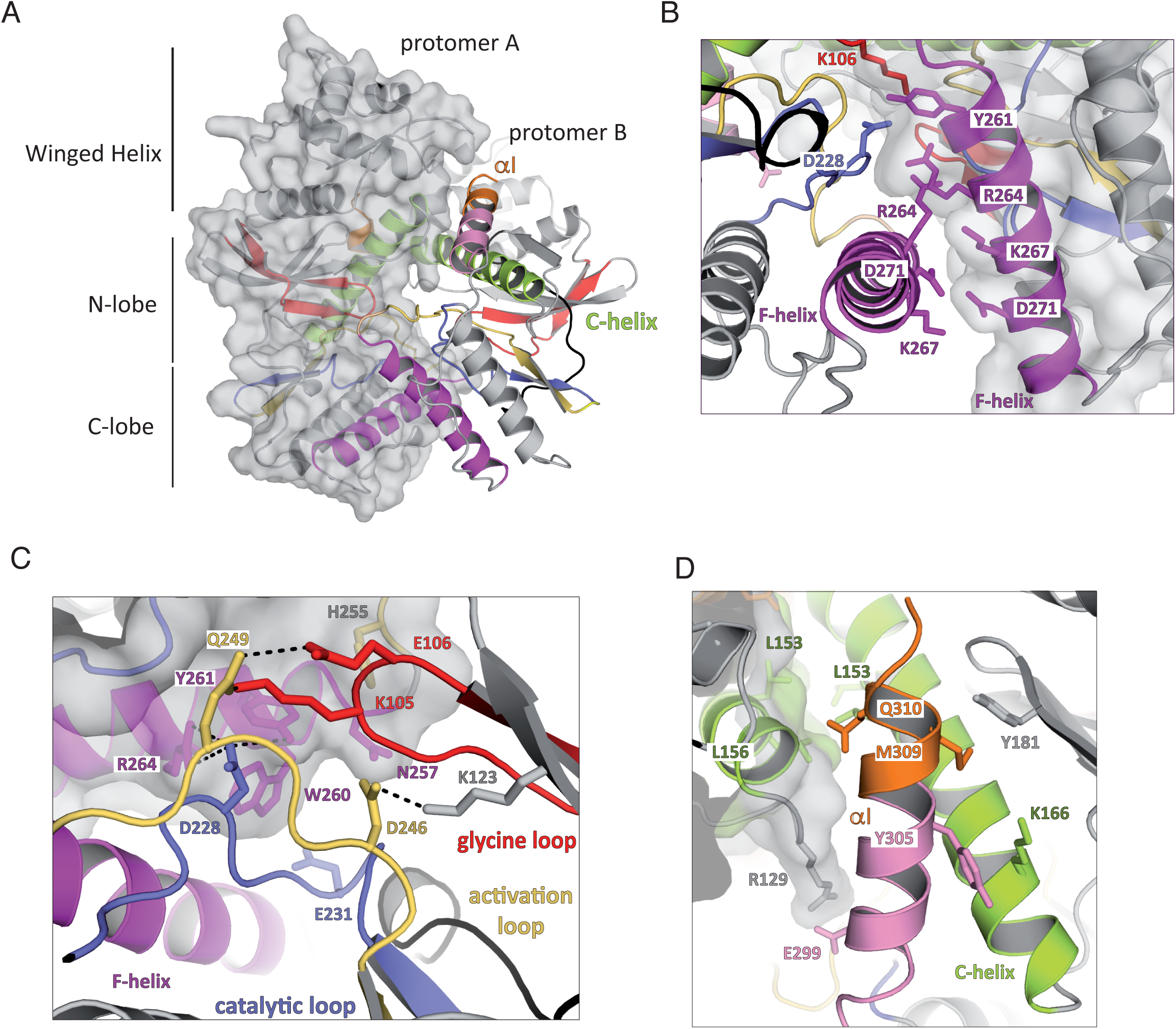
Human RIO2 forms homodimers. **A)** Human RIO2 interacts with itself to form an interconnected homodimer with a head to head orientation. A general view of hRIO2 homodimer is shown under the same orientation as in Fig. 1A with protomer A maintained in the same orientation. **B)** Close-up view of the homodimer interface around the F-Helix. One protomer is shown with a grey surface. Salt bridges between residues D271 and K267, and residues K267 and D271 are maintaining the two F-helices in close interaction. R264 of each protomer is stacking together through Van der Waals interactions, while Y261 is involved in a H-bond network with K106 and D228 of the other protomer. **C)** Close-up view of the homodimer interface around the ATP-binding region. One protomer is shown with a grey surface. The catalytic residue D228 is involved in H-bonding with Y261 and K106 while K105 backbone interacts with H255 and Q249 stabilizes K105 side chain. E231 is interacting with W260. The catalytic residues K123 and D246 are shown. **D)** Close-up view of the homodimer interface around C-Helix. The C-terminal *α*-helix is coloured in pink and orange. Aliphatic residues such as L153 and L156 are providing Van Der Waals interaction force that add to the many other interactions between the two protomers.

The ATP binding regions, via the glycine loop (residues K105 and E106), the activation loop (D246 and Q249) and the catalytic loop (D228 and E231), provide a significant portion of the interacting surface (Fig. 2C). Numerous polar interactions are found between D228 of one protomer and the hydroxyl groups of Y261 and R264 of the other protomer (Fig. 2C). Each C-helix of a protomer (light green) is facing the C-helix of the opposite protomer in an antiparallel mode (Fig. 2D), and this interaction is mainly restricted to Van der Waals contacts provided by L153 and L156. Altogether, the complete surface of interaction involves 1,570 Å^2^ per protomer.

In the longest crystallized form of hRIO2 (residues 1 to 353), some additional C-terminal residues form an *α−*helix packing at the interface between the two protomers to further extend the complex surface of interaction (residue E299 and R129; Fig. 2A and 2D). Importantly, all solved structures of hRIO2 show a dimeric organisation in the asymmetric unit. Taken together with the results from the sedimentation experiments, the observed dimerization of hRIO2 might mimic a biological situation.

### The *α*I of ctRio2 adopts a different conformation in hRIO2

Apart from the oligomeric state, a significant difference between hRIO2 and ctRio2 models is the position of the eukaryotic-specific *α*-helix (*α*I, orange), which packs in close vicinity to the ATP-binding site in ctRio2 ^26^(Fig. 3A and 3B). In ctRio2, the C-terminal residues (325-343) form a long *α*-helix (*α*I) that wraps around the protein and contributes to the ATP binding site environment (Fig. 3B). *α*I interacts with the N- and C-kinase lobes of the protein and is linked to the remainder of the protein through a short *α*-helix (residues 317-322, pink) embedded within a loop (Fig. 3B). In hRIO2, most of the equivalent residues form a continuous *α*-helix that packs against the C-helix, away from the ATP binding site (Fig. 3C). Due to hRIO2 homodimerization, *α*I cannot adopt the same orientation as observed in ctRio2. In the recent crystal structure of hRIO2 bound to an inhibitor ^33^, the same region of the protein adopts a different secondary structure forming a *β*-sheet with residues of the partially disordered *β*6-*α*5 loop (Fig. 3D). Even though deletion of the *β*6-*α*5 loop (residues 131-146) in our study does not alter the overall structure of hRIO2 or its oligomeric organization, the *β*6-*α*5 loop in hRIO2 (6HK6), partially occupies the position where the helical extension lies (this study). Thus the *α*-helix observed here is probably favoured by the deletion of the *β*6-*α*5 loop in our construct.

**Fig. 3:**
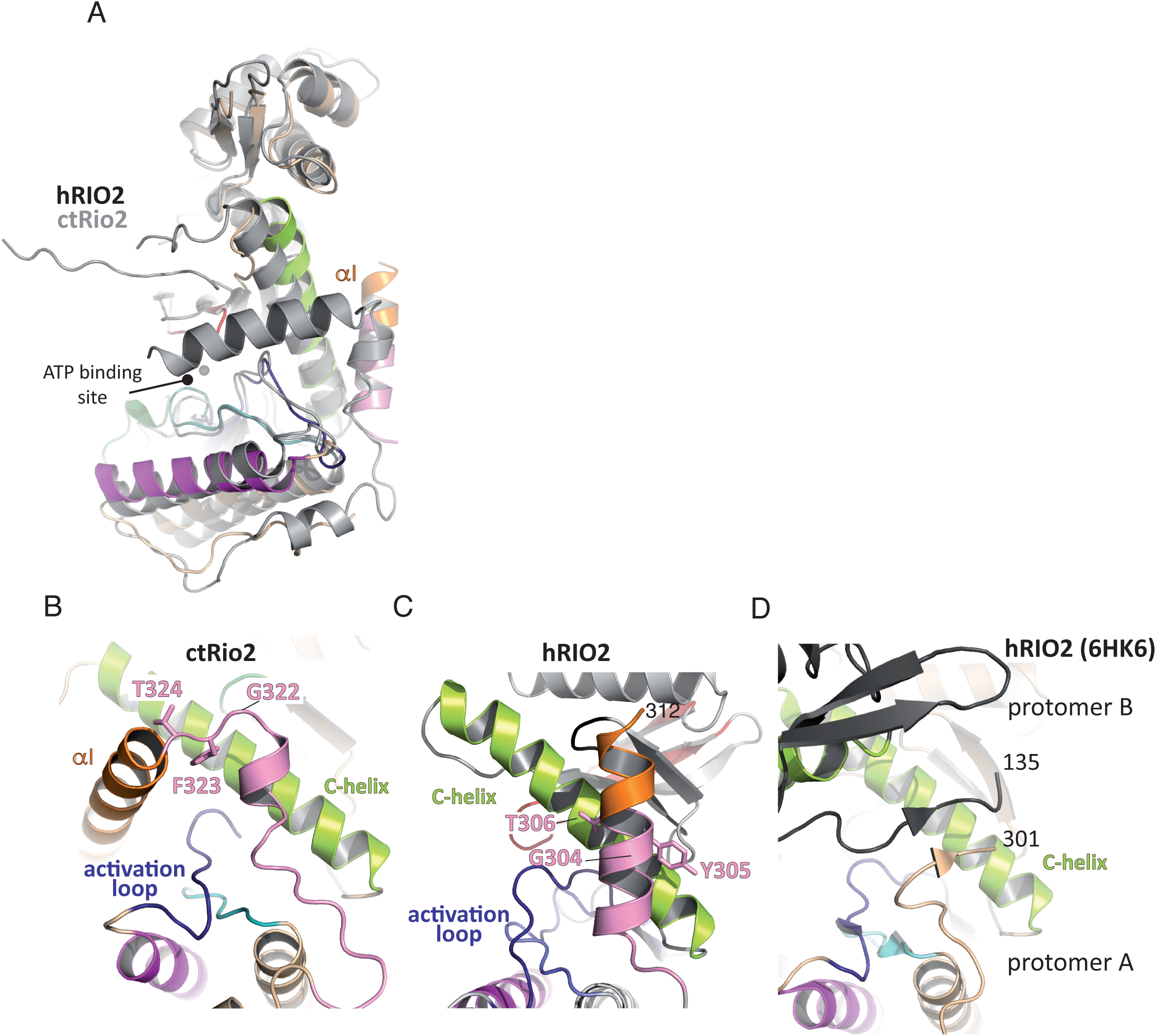
*α*I can adopt two conformational orientations. **A)** Superimposition of hRIO2(1-353)*Δ*(131-146) and ctRio2 has been performed in Coot ^59^. ctRio2 is displayed in grey. **B)** Focus on the *α*I of ctRio2. The residues 311 to 324 (pink and orange) of ctRio2 form a short *α*-helix embedded in a linker region that packs against the C-helix (light green). The GF/YT switch region is labelled. **C)** The similar region of hRIO2 corresponding to residues 311 to 324 of ctRio2 are displayed in the same orientation as in B. This region adopts an extended *α*-helix conformation that projects the C-terminal end of the downstream *α*-helix residues out of the ATP-binding pocket vicinity. **D)** The similar region of hRIO2 (PDB 6HK6) ^33^ is displayed under the same orientation as in B and C. The C-terminal residues of one protomer of hRIO2 form a *β*-sheet with residues from the *β*6-*α*5 loop of the other protomer (in grey).

### RIO2 homodimerization surface is conserved in eukaryotes

To the best of our knowledge, dimerization of afRio2 or ctRio2 has not been observed in the published crystal structures ^26,30^. In an attempt to understand this discrepancy with hRIO2, we looked at sequence alignments of Rio2 protein orthologues to identify hot spots of residue conservation (Fig. S3). We then plotted the residue conservation across species on the surface of hRIO2 atomic model using the webserver ConSurf ^34^ (Fig. 4). When all species are considered, sequence conservation is limited to the ATP-binding pocket (Fig. 4A and 4B). However, when sequence conservation is calculated using only eukaryotic sequences or multicellular eukaryotic sequences, invariant residues cluster not only at the ATP-binding pocket but also along the dimer interface (Fig. 4C to 4E). Of note, the same surface area is used for the association of Rio2 with 20S pre-rRNA (helices 1, 18, 28, 34, and 44), Tsr1, Ltv1, uS12, us7, uS13 and uS19 ^28,29^. The overlapping mechanism for a subset of residues that are involved in pre-40S particle association and dimer formation suggests that if dimerization exist *in vivo*, it is incompatible with pre-40S association. The superimposition of ctRio2 and hRIO2 suggests that *α*I prevents ctRio2 dimer formation. This observation is in agreement with the absence of the ctRio2 dimer in AUC experiments. It is also interesting to note that the eukaryotic-specific *α*I helix is disordered in the recent structure of the pre-40S particle ^28,29^, suggesting that the *α*I is a mobile secondary structure that may not be involved in ATP catalysis in pre-40S maturation.

**Fig. 4:**
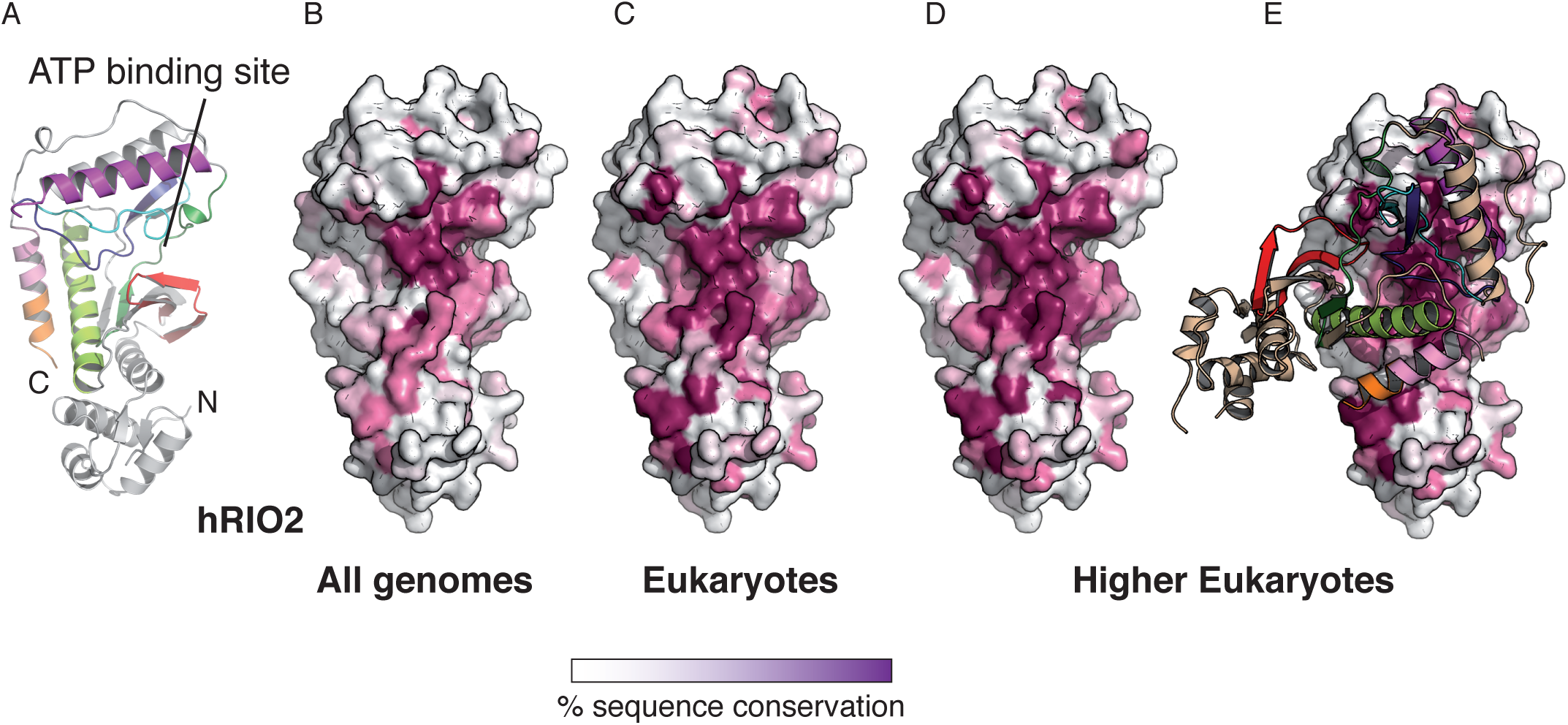
RIO2 homodimer interface is conserved in eukaryotes. The sequence conservation was calculated using the Consurf server (http://consurf.tau.ac.il/) and the sequence alignment is displayed in Fig. S2. The conservation above 70 %, 80 %, 90 % and equal to 100 % are shown in a gradient from white to magenta. **A)** Structure of hRIO2(1-353)*Δ*(131-146) protomer. The N- and C-termini are labelled, as well as the ATP-binding pocket. **B)** Sequence conservation in all Rio2 orthologues is displayed at the surface of hRIO2 and cluster at the ATP-binding pocket. **C)** Sequence conservation in eukaryotic Rio2 protein sequences is displayed at the surface of hRIO2. The fingerprint extends around the ATP-binding pocket. **D) and E)** Sequence conservation in higher eukaryotes (*Homo sapiens*, *Xenopus laevis*, *Drosophila melanogaster*, *Caenorhabditis elegans*, *Zea mais*, *Dictyostelium discoideum*, *Gallus gallus*, *Aedes aegypti*, *Danio rerio*) is displayed at the surface of hRIO2 structure. The region of conservation covers the entire homodimer interface. The bar indicates the residue conservation from white (non-conserved) to magenta (identical residues).

### Homodimerization alters the ATP-binding pocket

The afRio2 and ctRio2 structures have been solved in complex with various ligands including ADP, ATP and toyocamycin ^14,26^. In order to analyse the local rearrangements of the C*α* backbone around the ATP binding site, the ADP-bound structure of ctRio2 and the hRIO2 structure were superimposed and the overall structure around the ADP is shown (Fig. 5). In comparison to the ADP-bound structure of ctRio2, the ligand-binding pocket of hRIO2 has undergone substantial local rearrangements. The main difference occurs at the glycine loop (red) that adopts a closed conformation in hRIO2. In this orientation, the glycine loop prevents ATP binding by sterical hindrance at a position where the *γ*-phosphate of an ATP molecule would sit (Fig. 5A and 5B). However, even though full ATP cannot be accommodated in that conformation, a smaller molecule such as an inhibitor can fit as demonstrated by the crystal structure of hRIO2 in complex with compound 9 ^33^. Of note, this inhibitor occupies a position in the ligand binding pocket where the adenine moiety is recognized. The glycine loop of one protomer is stabilized by a series of interactions with residues D228, R264 and Y261 of the adjacent protomer (Fig. 5B, grey shading) while the glycine loop is also stabilized through interactions between G104 and H255, E106 and Q249. Remarkably every residue involved in the glycine loop stabilization is highly conserved across species (Fig. S3). Despite many attempts to soak crystals with various ATP analogues, or to incubate the protein sample prior crystallization with ATP analogues, we never managed to obtain a liganded form of hRIO2 nor managed to crystallize a monomeric from. Altogether, these findings suggest a concerted molecular mechanism by which hRIO2 homodimerization favours a non-liganded form of the protein.

**Fig. 5:**
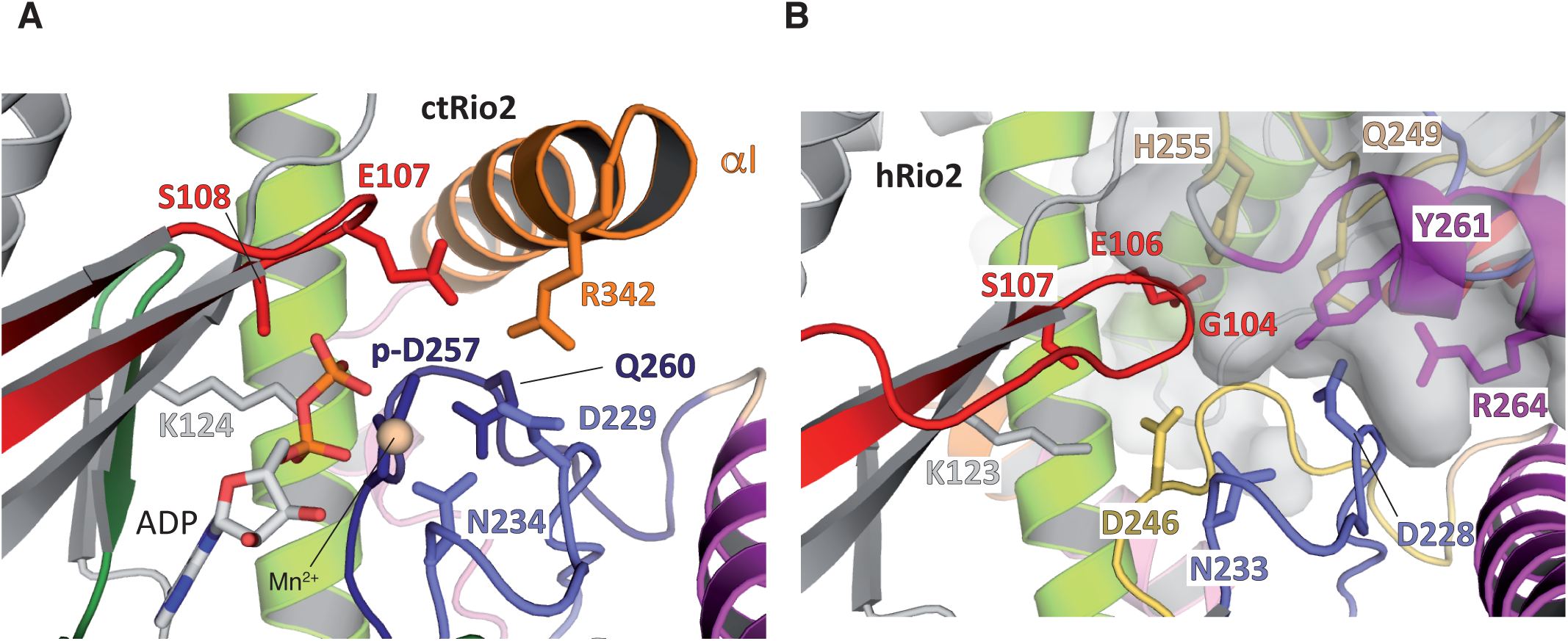
Homodimer formation closes the ATP-binding pocket. ctRio2 and hRIO2 structures have been superimposed and are displayed separately. **A)** The ATP-binding pocket of ctRio2 in complex with ADP (PDB code 4GYI) ^26^. Residues involved in the ADP binding and the phospho-aspartate residue (p-D257) are indicated. A Mn^2+^ ion is also shown in beige. **B)** Human RIO2 ATP-binding pocket is shown under same orientation as ctRio2. The equivalent residues involved in ADP binding in ctRio2 are displayed. Residues belonging to the adjacent hRIO2 protomer are shaded in grey.

## DISCUSSION

In this article, we report the crystal structure of three different constructs of human RIO2 protein kinase. The structures extend from the N-terminus up to the residues 290 and 312 out of 552 residues of full length hRIO2. Homodimeric forms of hRIO2 were observed in all cases, and no ligand was detected in the ATP-binding pocket. Dimer formation relies on a network of interactions resulting in the inward movement of the glycine loop towards the ATP-binding pocket leading to the ATP-binding pocket partial closure. The glycine loop (P-loop) as a role in ATP hydrolysis stimulation and pre-40S binding ^27^. Residues involved in dimer formation are highly conserved in higher eukaryotes. However, these same residues are also involved in pre-40S particle interaction. In the absence of *in vivo* evidence, we cannot exclude the possibility that these residues are conserved only for pre-40S particle association.

Recently, another group reported the crystal structure of hRIO2 in complex with an inhibitor ^33^. In their study, a C-terminally truncated version (residues 1 to 329) of hRIO2 crystallized with 10 molecules per asymmetric unit among which only eight are found as dimers and two molecules as protomers. This observation, in addition to a SEC analysis, suggested that hRIO2 is a monomer in solution. Inspection of the crystal lattice shows that the two isolated protomers found in an asymmetric unit form dimers with neighbouring protomers in the adjacent asymmetric unit. Thus, no isolated protomer is really observed in the crystal structure. As we also observe, dimerization of hRIO2 increases with protein concentration but independently of the presence of an inhibitor. The inhibitor observed in Wang et al. lies at the interface between two molecules in the crystal packing and not at the dimer interface *per se*. Thus, the inhibitor is likely to be essential for protein crystallization in their conditions but not for dimerization.

In other protein kinases, dimerization is a common feature driving their activation ^35,36^. For example, this process is described for EGFR, RAF and CHK2 kinases ^37^. In the case of B-RAF, dimerization occurs at the opposite face of the ATP-binding pocket and leaves the protein able to bind ATP ^38^. In general, the activation is thought to occur via an allosteric mechanism in which one of the protomer acts as a scaffolding protein (the activating kinase) to stabilize the active conformation of the other protomer (the receiving kinase). CHK2 protein kinase dimerization is promoted by the phosphorylation of residue T68 by the ATM kinase, which leads to CHK2 activation through auto-phosphorylation ^37^. Once phosphorylated the CHK2 dimer rapidly dissociates into protomers. In hRIO2, the dimer blocks the ATP-binding pockets in a conformation where ATP cannot be accommodated, in an opposite mechanism to CHK2. Another example of homodimer formation is the *Legionella pneumophila* LegK4 kinase ^39^. In the LegK4 structure, the protein dimerizes through *α*F, *α*G and the *α*G-*α*I loop of the C-lobe in both the apo and AMP-PNP bound states, which differs from the dimer formation observed for hRIO2. Finally, In the cryo-EM reconstructions of yeast and human pre-40S particles, only a protomer of Rio2 protein could be fit into the attributed density for Rio2 ^26,28,29,40,41^. Altogether, these data strongly suggest that the *in vitro* dimeric form of RIO2 is an inactive form.

The C-terminal residues described in ctRio2 and hRIO2 structures display different conformations. In the dimeric form of hRIO2, they provide additional interactions and extend the dimer interface. In the monomeric form of ctRio2, the C-terminal helix is incompatible with dimer formation and participates in interactions around the catalytic centre which might include an auto-inhibitory action toward ATP catalysis ^26^. Finally, in the pre-40S particle cryo-EM reconstruction, this eukaryotic specific *α*-helix is not observed ^28,29^.

Several point mutations impair hRIO2 kinase activity. This is the case for the K123A/D246A double mutation in hRIO2 ^13^ and the D229A substitution in Rio2 ^22^. On the basis of the ctRio2/ADP complex structures, the K123A/D246A double mutation affects the ATP recognition (Fig. 5A & 5B), while the D228A substitution may have a double effect on both the catalytic activity and the dimer formation. Indeed, in our structures the D228 side chain (D229 in ctRio2) points towards the neighbouring hRIO2 protomer and not towards the ATP binding pocket. This suggests that an identical set of residues is used for ATP recognition/catalysis, or peptide substrate activation and dimer formation. This would be an elegant molecular mechanism for segregating active versus inactive forms of the enzyme.

At this stage and in the absence of *in vivo* evidence for hRIO2 dimerization, the actual biological role of the hRIO2 dimer remains elusive and only speculations can be formulated. Efforts have been made to understand ribosomal protein and RNA trafficking across the nuclear pore complex ^42–44^, however there is a limited knowledge concerning the AFs required for ribosome biogenesis that have to be (re)imported to the nucleolus ^43,45^. For the yeast Rio2, it has been shown that it possesses a C-terminal NES required for the Crm1-driven pre-40S particle export ^32^. In human, hCRM1 binds residues 391 to 403 of hRIO2 as a NES ^31^. hRIO2 is found at low concentration in the cell in comparison to other AFs such as hNOB1 and hPNO1 ^25,46^. While being mainly cytosolic, it is incorporated at nuclear stages of pre-ribosome maturation. This implies an efficient import mechanism of recycling of hRIO2 as its quantity is a limiting step. Indeed, depletion of the free pool of Rio2 is observed upon prevention of its release from the pre-40S particles ^26^. The pool of available Rio2 is therefore fully immobilized on pre-40S particles. Having Rio2 as a dimer in its apo form may combine several advantages.

Dimerization would i) prevent re-association of Rio2 to pre-mature or mature 40S particles in the cytoplasm after its release, ii) prevent ATPase activity when not bound to the pre-40S particle, iii) be required for its import. For this latter point, no classical NLS can be identified from the primary sequence, and no importin has yet been linked to Rio2 import. Alternatively, the use of any importin in an opportunistic and serendipity manner could be envisaged. Dimer formation for nuclear import has been reported in the case of uS3 (RpS3). In this mechanism, the nuclear import of uS3 is chaperoned by Yar1 and the ternary complex enters the nucleus by Kap60/Kap95 ^43,47^. Dimer dissociation would have to be promoted by an external and yet unidentified factor. It has been shown in the case afRio1 and hRIO1 that the oligomeric state of the protein impacts on its kinase activity ^48^ and that auto-phosphorylation plays an important role in such a mechanism. The switch from a cytoplasmic inactive and dimeric state to a nuclear active and monomeric state remains further study. However, it provides an exciting regulatory mechanism for specific AF re-import to the nucleus after their cytoplasmic release.

## MATERIALS AND METHODS

### Constructs

All constructs were sequenced to ensure the absence of mutations. The human RIO2 kinase constructs 1-321, as well as 1-353 and 1-321 with residues 131 to 150 deleted, were amplified from the DNA encoding the full-length hRIO2 and subcloned into the *Nde*I and *BamH*I sites of a modified pET-15b (Novagen) plasmid ^49^ to produce an N-terminal His-tag fused protein containing a TEV cleavage site and an ampicillin resistance marker. The *Chaetomium thermophilum* (ct) Rio2 (kind gift from N. Laronde-Leblanc) was subcloned into a pET-15b derived plasmid to generate an N-terminal His-tagged protein.

### Protein expression and purification

The various constructs were transformed in *E. coli* BL21(DE3) Gold cells. The cultures were grown at 37 °C in 2xYT medium supplemented with 100 mg/L ampicillin and 30 mg/L kanamycin. Cells were induced at 20 °C with 0.5 mM IPTG for 14 hours and collected by centrifugation at 4,500 g and resuspended in buffer containing 50 mM Tris pH 8.0, with 350 mM or 900 mM NaCl. Cell pellets were lysed with an Emulsi Flex-C3 (Avestin) and centrifuged at 50,000 g. The cell lysate was combined with His-Select Co^2+^ resin (Sigma) for 1 hour at 4 °C. The resin containing bound proteins was washed with 5 column volumes of resuspension buffer and the protein complexes were eluted with a 10 mM - 250 mM imidazole gradient. hRIO2 (1-321), hRIO2 (1-321)*Δ*(131-146), and hRIO2 (1-353) *Δ*(131-146) were further purified with a HiQ-Sepharose (GE Healthcare) chromatography, followed by size exclusion chromatography over a Superdex 200 column (GE Healthcare). The selenomethionine substituted hRIO2 (1-321) *Δ* (131-146) protein was purified in the same manner as native hRIO2 (1-321).

### Analytical Ultracentrifugation

Analytical ultracentrifugation experiments were performed in a Beckman Coulter ProteomeLab XL-I instrument (IGBMC Structural Biology Platform) at 4°C. ctRio2 and hRIO2(1-321)*Δ*(131-46) proteins were independently purified and dialyzed against a buffer containing 25 mM Tris-HCl pH 7.4, 300 mM NaCl and 5% Glycerol (w/v). For analytical ultracentrifugation sedimentation velocity experiments at 50,000 RPM, 400 µl of a concentration series of each protein was prepared into the dialysis buffer at a final concentration of 60, 30, 15, 7.5 and 3.75 µM for ctRio2 and 54, 27, 13.5, 6.75 and 3.375 µM for hRIO2. Absorbance scans at 280 nm and interference scans were taken every 5 minutes. Sedimentation data were first analyzed using SEDFIT software ^50^ and the continuous sedimentation coefficient distribution model c(s) were generated and then exported into SEDPHAT for fitting with the hybrid local continuous distribution model ^51^. Buffer density, buffer viscosity and protein partial specific volumes were calculated using SEDNTERP software. GUSSI was used to integrate the sedimentation peaks and to produce the graphs ^52^.

### Crystallization & Structure determination

Prior to crystallization, the native hRIO2 (1-321), hRIO2 (1-353)*Δ*(131-146) and the selenomethionine substituted hRIO2 (1-321)*Δ*(131-146) proteins were exchanged in buffer containing 25 mM Tris, pH 7.5, 900mM or 350 mM NaCl (respectively), 1.0 mM DTT and concentrated to 10 mg/ml. The complexes were crystallized at 20°C by the sitting drop method against a reservoir containing condition D2 or G2 of the Morpheus™ screen (Molecular Dimensions). Crystals were transferred to a cryo protectant with 60 % EDO-P8K, flash-frozen in liquid nitrogen, and maintained at 100 K with a nitrogen cryostream during data collection.

Crystals of hRIO2 (1-321) and hRIO2 (1-321)*Δ*(131-146) belong to the *P*2_1_ space group with unit cell dimensions a=69.13 Å b=88.30 Å c=102.14 Å and contain four molecules per asymmetric unit. Crystals of hRIO2 (1-353)*Δ*(131-146) belong to the *P*6_5_ space group with unit cell dimensions a=67.55Å b=67.55 Å c=312.2 Å

The structure of the hRIO2 (1-321)*Δ*(131-146) was solved by SAD using selenomethionine-substituted protein crystals. Datasets were reduced using XDS ^53^. 40 selenium sites were located with SHELXCD ^54^ and refined with Phaser ^55^. The initial model was build using Bucaneer ^56^. The final models were refined with BUSTER 2.10 ^57^ and display good stereochemistry (Table 1). The atomic model of hRIO2 (1-321)*Δ*(131-146) comprises residues 6-312. The atomic model of hRIO2 (1-353)*Δ*(131-146) comprises residues 9-314. The atomic model of hRIO2 (1-321) comprises residues 12 to 291 with the exception of loops 66-70 and 129-149.

## ACKNOWLEDGEMENTS

We acknowledge the European Synchrotron Radiation Facility for provision of synchrotron radiation facilities and we would like to thank staff members for assistance in using beamline ID14-1 and ID29. We also acknowledge the Synchrotron SOLEIL and Drs Pierre Legrand and Andrew Thompson for providing access to Proxima-1 beamline. We acknowledge N. Laronde-Leblanc for providing the ctRio2 plasmid. We thank C.D. Mackereth for critical reading of the manuscript.

Molecular graphics and analyses were performed with the UCSF Chimera package. Chimera is developed by the Resource for Biocomputing, Visualization, and Informatics at the University of California, San Francisco (supported by NIGMS P41-GM103311).

## FUNDING

This work was supported by an ANR Blanc 2010 grant RIBOPRE40S, University of Bordeaux, CNRS and INSERM. The authors acknowledge the support and the use of resources of the French Infrastructure for Integrated Structural Biology FRISBI ANR-10-INBS-05 and of Instruct-ERIC.

## AUTHORS CONTRIBUTION

FM and NP expressed, purified and crystallized human Rio2. ST performed the kinase assays. SF solved and refined the structure. SF analyzed the data, wrote the paper and supervised the work.

## DISCLOSURE STATEMENT

The authors declare that they have no competing financial interests.

## Supporting information

### Supporting Materials and Methods

#### Kinase assay

The autophosphorylation activity of RIO2 kinase was tested using the Rio2 protein from *Chaetomium thermophilum* and from *Homo sapiens*. For the later species, three constructs were assayed: the full-length protein (hRIO2), the truncation corresponding to res. 1 to 353 including loop deletion between res. 131 to 146 (hRIO2(1-353)Δ(131-146)) and the truncation from res. 1 to 321 including loop deletion between res. 131 to 146 (hRIO2(1-321)Δ131-146). The proteins were purified to homogeneity using the protocol indicated in the Materials and Methods section. Each protein was incubated with γ-[32P] ATP for 120 min at 37 °C in a buffer containing 25 mM Tris pH 7.4, 150 mM KCl, 10 % Glycerol and 5 mM MgCl_2_. Control reactions were incubated with additional 50 mM EDTA to chelate magnesium ions. All the reactions were stopped by the addition of Laemmli buffer and loaded on a 12.5 % SDS-PAGE, ran for 2 h at 160 V and autoradiographed.

**Figure S1:**
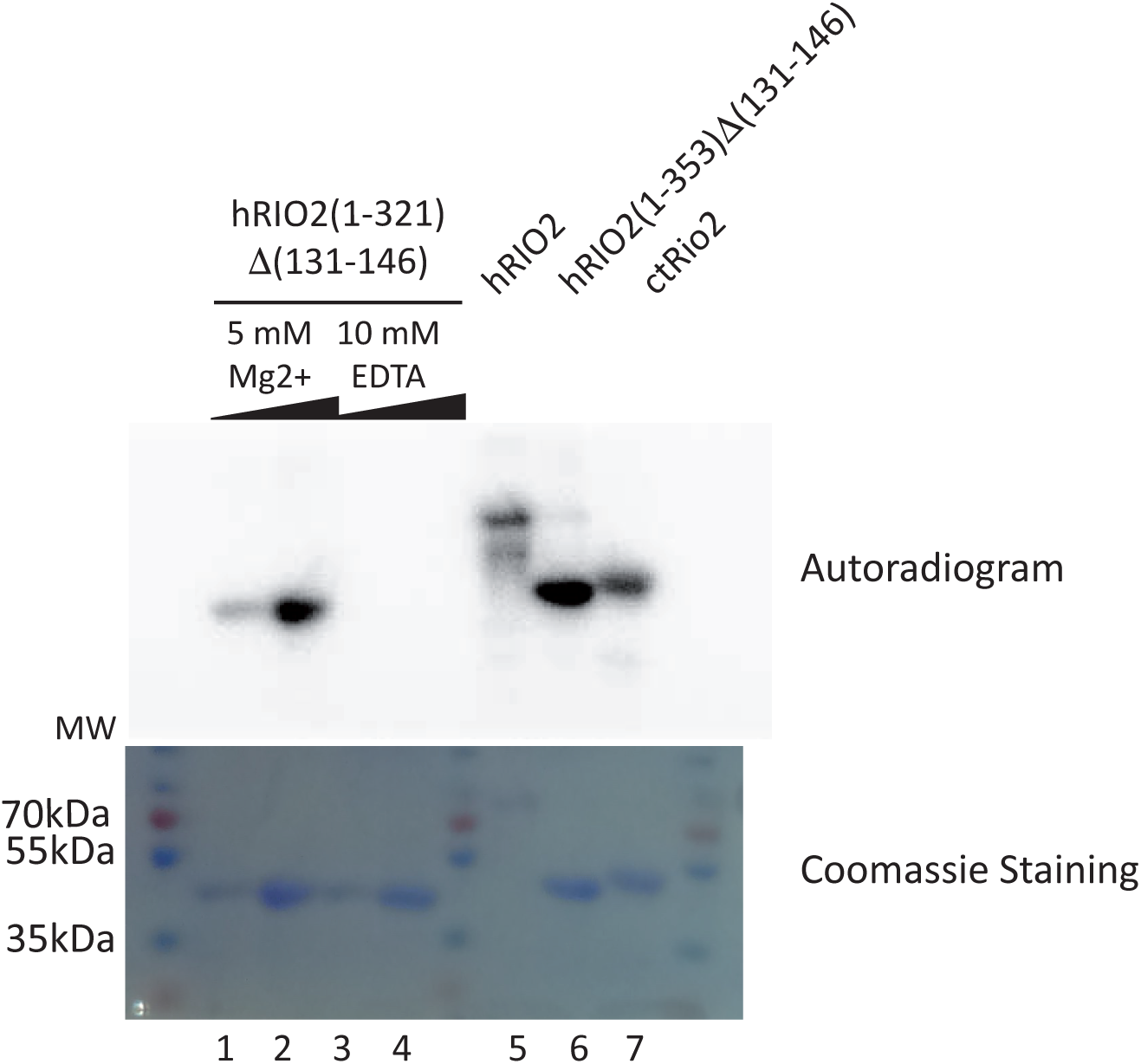
hRIO2 autophosphorylation activity. The autophosphorylation activity test of Rio2 kinase is described in the supplementary data. The hRIO2(1-321)Δ(131-146) was tested using two amounts of protein (3 µg and 12 µg, lane 1 and 2 respectively). Control reactions were measured in presence (lane 1 & 2) or absence (Lane 3 & 4) of Mg^2+^. Three other constructs were tested and used as positive controls (hRIO2 full length and ctRio2) for hRIO2(1-353)Δ(131-146). Each experiment was performed with 10 µg of protein under the same experimental condition. The autophosphorylation activity of Rio2 was assayed as described in Zemp et al. ^1^. hRIO2 full-length, ctRio2 were used as positive controls for the hRIO2(1-321)Δ(131-146) and hRIO2(1-353)Δ(131-146).

**Figure S2:**
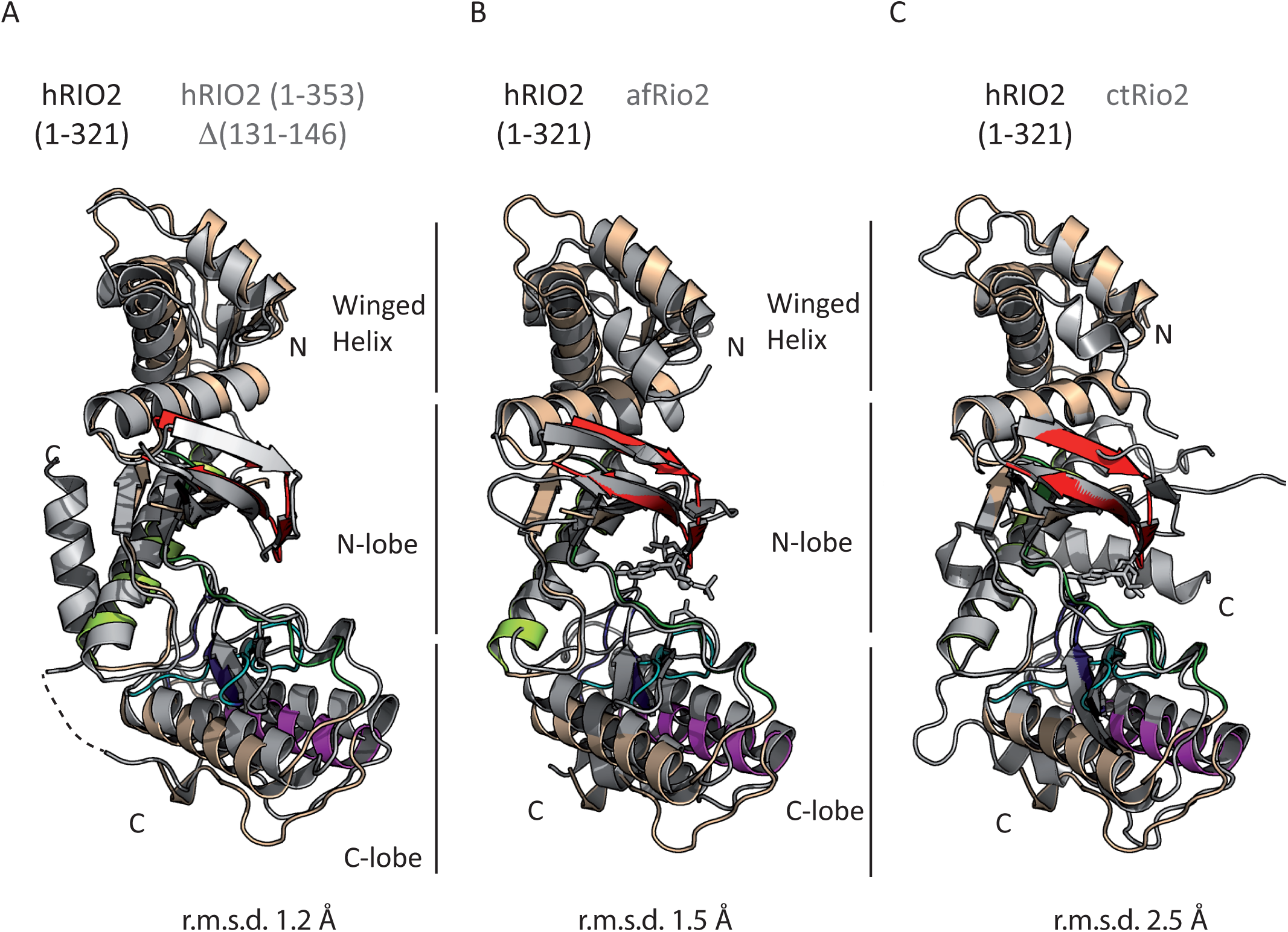
Overall similarity of hRIO2, afRio2 and ctRio2. The human RIO2(1-321) model was used as a reference to superimpose hRIO2(1-321)Δ(131-146)(A), afRio2 (B) and ctRio2 (C) shown in grey. The overall r.m.s.d. is indicated beneath the figure. The prototypical kinase motifs are represented in colour. The so-called C-helix (light green), F-helix (magenta), glycine loop (red), catalytic loop (blue) and activation loop (black) are displayed.

**Figure S3:**
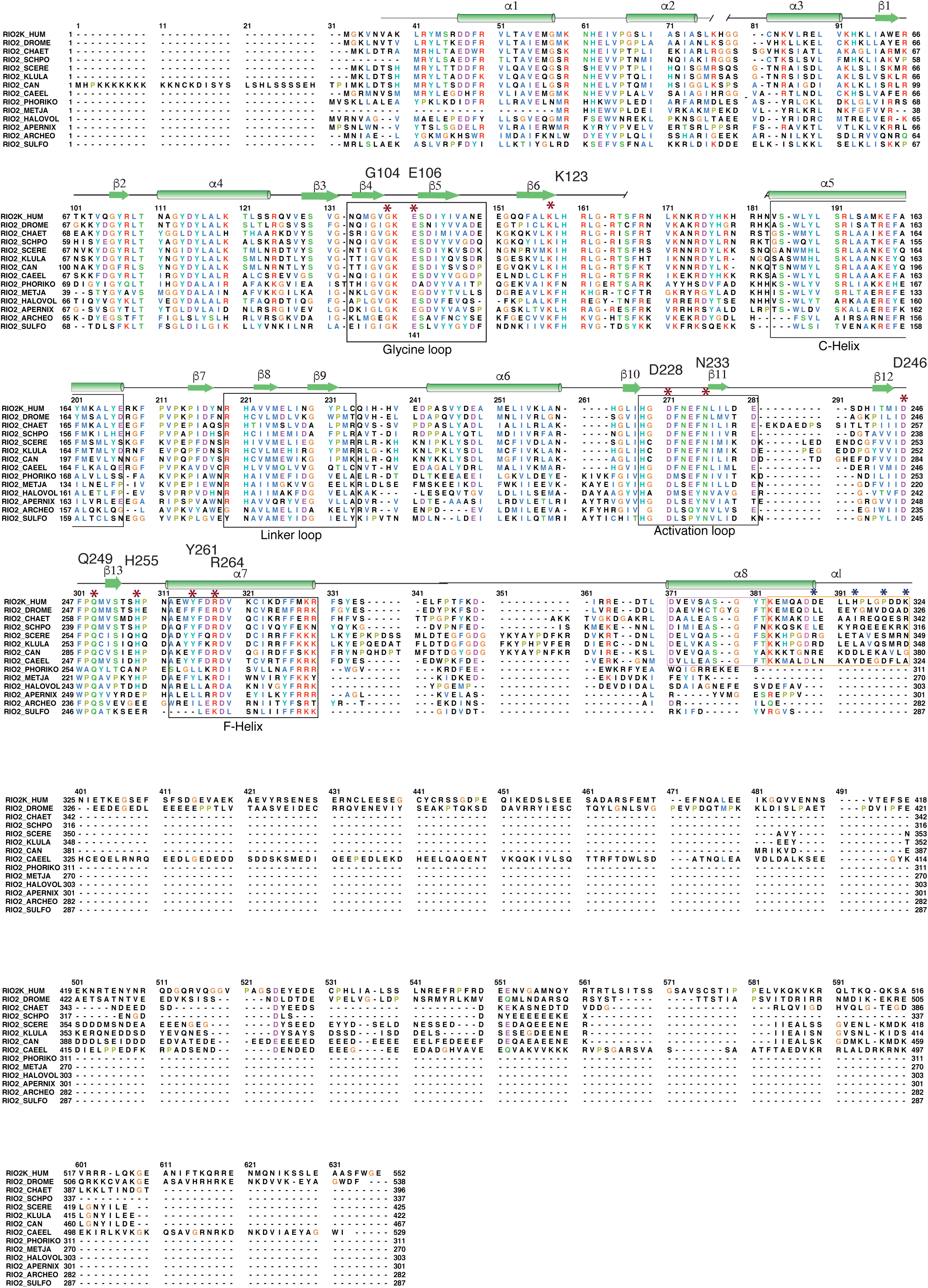
Sequence alignment of Rio2 orthologues. Sequence alignment has been performed using Chimera ^2^ and displayed with the help of SEAVIEW ^3^. The secondary structures are displayed on top of the sequence. Key residues discussed in the manuscript are labelled with a red star.

## REFERENCES

1. Henras AK, Plisson-Chastang C, O’Donohue MF, Chakraborty A, Gleizes PE. An overview of pre-ribosomal RNA processing in eukaryotes. Wiley Interdiscip Rev RNA [Internet] 2015; 6:225–42. Available from: http://www.ncbi.nlm.nih.gov/pubmed/25346433

2. de la Cruz J, Karbstein K, Woolford Jr. JL. Functions of ribosomal proteins in assembly of eukaryotic ribosomes in vivo. Annu Rev Biochem [Internet] 2015; 84:93–129. Available from: http://www.ncbi.nlm.nih.gov/pubmed/25706898

3. Bassler J, Grandi P, Gadal O, Lessmann T, Petfalski E, Tollervey D, Lechner J, Hurt E. Identification of a 60S preribosomal particle that is closely linked to nuclear export. Mol Cell [Internet] 2001; 8:517–29. Available from: http://www.ncbi.nlm.nih.gov/entrez/query.fcgi?cmd=Retrieve&db=PubMed&dopt=Citation&list_uids=11583615

4. Dragon F, Gallagher JE, Compagnone-Post PA, Mitchell BM, Porwancher KA, Wehner KA, Wormsley S, Settlage RE, Shabanowitz J, Osheim Y, et al. A large nucleolar U3 ribonucleoprotein required for 18S ribosomal RNA biogenesis. Nature [Internet] 2002; 417:967–70. Available from: http://www.ncbi.nlm.nih.gov/entrez/query.fcgi?cmd=Retrieve&db=PubMed&dopt=Citation&list_uids=12068309

5. Harnpicharnchai P, Jakovljevic J, Horsey E, Miles T, Roman J, Rout M, Meagher D, Imai B, Guo Y, Brame CJ, et al. Composition and functional characterization of yeast 66S ribosome assembly intermediates. Mol Cell [Internet] 2001; 8:505–15. Available from: http://www.ncbi.nlm.nih.gov/entrez/query.fcgi?cmd=Retrieve&db=PubMed&dopt=Citation&list_uids=11583614

6. Milkereit P, Kuhn H, Gas N, Tschochner H. The pre-ribosomal network. Nucleic Acids Res [Internet] 2003; 31:799–804. Available from: http://www.ncbi.nlm.nih.gov/entrez/query.fcgi?cmd=Retrieve&db=PubMed&dopt=Citation&list_uids=12560474

7. Nissan TA, Bassler J, Petfalski E, Tollervey D, Hurt E. 60S pre-ribosome formation viewed from assembly in the nucleolus until export to the cytoplasm. Embo J [Internet] 2002; 21:5539–47. Available from: http://www.ncbi.nlm.nih.gov/entrez/query.fcgi?cmd=Retrieve&db=PubMed&dopt=Citation&list_uids=12374754

8. Schafer T, Strauss D, Petfalski E, Tollervey D, Hurt E. The path from nucleolar 90S to cytoplasmic 40S pre-ribosomes. Embo J [Internet] 2003; 22:1370–80. Available from: http://www.ncbi.nlm.nih.gov/entrez/query.fcgi?cmd=Retrieve&db=PubMed&dopt=Citation&list_uids=12628929

9. Rouquette J, Choesmel V, Gleizes PE. Nuclear export and cytoplasmic processing of precursors to the 40S ribosomal subunits in mammalian cells. Embo J [Internet] 2005; 24:2862–72. Available from: http://www.ncbi.nlm.nih.gov/entrez/query.fcgi?cmd=Retrieve&db=PubMed&dopt=Citation&list_uids=16037817

10. Baumas K, Soudet J, Caizergues-Ferrer M, Faubladier M, Henry Y, Mougin A. Human RioK3 is a novel component of cytoplasmic pre-40S pre-ribosomal particles. RNA Biol [Internet] 2012; 9:162–74. Available from: http://www.ncbi.nlm.nih.gov/entrez/query.fcgi?cmd=Retrieve&db=PubMed&dopt=Citation&list_uids=22418843

11. Widmann B, Wandrey F, Badertscher L, Wyler E, Pfannstiel J, Zemp I, Kutay U. The kinase activity of human Rio1 is required for final steps of cytoplasmic maturation of 40S subunits. Mol Biol Cell [Internet] 2012; 23:22–35. Available from: http://www.ncbi.nlm.nih.gov/entrez/query.fcgi?cmd=Retrieve&db=PubMed&dopt=Citation&list_uids=22072790

12. Ferreira-Cerca S, Kiburu I, Thomson E, Laronde N, Hurt E. Dominant Rio1 kinase/ATPase catalytic mutant induces trapping of late pre-40S biogenesis factors in 80S-like ribosomes. Nucleic Acids Res 2014; 42:8635–47.

13. Zemp I, Wild T, O’Donohue MF, Wandrey F, Widmann B, Gleizes PE, Kutay U. Distinct cytoplasmic maturation steps of 40S ribosomal subunit precursors require hRio2. J Cell Biol [Internet] 2009; 185:1167–80. Available from: http://www.ncbi.nlm.nih.gov/entrez/query.fcgi?cmd=Retrieve&db=PubMed&dopt=Citation&list_uids=19564402

14. LaRonde-LeBlanc N, Guszczynski T, Copeland T, Wlodawer A. Autophosphorylation of Archaeoglobus fulgidus Rio2 and crystal structures of its nucleotide-metal ion complexes. Febs J [Internet] 2005; 272:2800–10. Available from: http://www.ncbi.nlm.nih.gov/entrez/query.fcgi?cmd=Retrieve&db=PubMed&dopt=Citation&list_uids=15943813

15. Endicott JA, Noble ME, Johnson LN. The structural basis for control of eukaryotic protein kinases. Annu Rev Biochem [Internet] 2012; 81:587–613. Available from: http://www.ncbi.nlm.nih.gov/pubmed/22482904

16. Vanrobays E, Gleizes PE, Bousquet-Antonelli C, Noaillac-Depeyre J, Caizergues-Ferrer M, Gelugne JP. Processing of 20S pre-rRNA to 18S ribosomal RNA in yeast requires Rrp10p, an essential non-ribosomal cytoplasmic protein. Embo J [Internet] 2001; 20:4204–13. Available from: http://www.ncbi.nlm.nih.gov/entrez/query.fcgi?cmd=Retrieve&db=PubMed&dopt=Citation&list_uids=11483523

17. Vanrobays E, Gelugne JP, Gleizes PE, Caizergues-Ferrer M. Late cytoplasmic maturation of the small ribosomal subunit requires RIO proteins in Saccharomyces cerevisiae. Mol Cell Biol [Internet] 2003; 23:2083–95. Available from: http://www.ncbi.nlm.nih.gov/entrez/query.fcgi?cmd=Retrieve&db=PubMed&dopt=Citation&list_uids=12612080

18. Iacovella MG, Golfieri C, Massari LF, Busnelli S, Pagliuca C, Dal Maschio M, Infantino V, Visintin R, Mechtler K, Ferreira-Cerca S, et al. Rio1 promotes rDNA stability and downregulates RNA polymerase I to ensure rDNA segregation. Nat Commun [Internet] 2015; 6:6643. Available from: http://www.ncbi.nlm.nih.gov/pubmed/25851096

19. Angermayr M, Roidl A, Bandlow W. Yeast Rio1p is the founding member of a novel subfamily of protein serine kinases involved in the control of cell cycle progression. Mol Microbiol [Internet] 2002; 44:309–24. Available from: http://www.ncbi.nlm.nih.gov/entrez/query.fcgi?cmd=Retrieve&db=PubMed&dopt=Citation&list_uids=11972772

20. Angermayr M, Hochleitner E, Lottspeich F, Bandlow W. Protein kinase CK2 activates the atypical Rio1p kinase and promotes its cell-cycle phase-dependent degradation in yeast. Febs J [Internet] 2007; 274:4654–67. Available from: http://www.ncbi.nlm.nih.gov/entrez/query.fcgi?cmd=Retrieve&db=PubMed&dopt=Citation&list_uids=17725716

21. Leger-Silvestre I, Milkereit P, Ferreira-Cerca S, Saveanu C, Rousselle JC, Choesmel V, Guinefoleau C, Gas N, Gleizes PE. The ribosomal protein Rps15p is required for nuclear exit of the 40S subunit precursors in yeast. Embo J [Internet] 2004; 23:2336–47. Available from: http://www.ncbi.nlm.nih.gov/entrez/query.fcgi?cmd=Retrieve&db=PubMed&dopt=Citation&list_uids=15167894

22. Geerlings TH, Faber AW, Bister MD, Vos JC, Raue HA. Rio2p, an evolutionarily conserved, low abundant protein kinase essential for processing of 20 S Pre-rRNA in Saccharomyces cerevisiae. J Biol Chem [Internet] 2003; 278:22537–45. Available from: http://www.ncbi.nlm.nih.gov/entrez/query.fcgi?cmd=Retrieve&db=PubMed&dopt=Citation&list_uids=12690111

23. Sloan KE, Knox A, Wells G, Schneider C, Watkins NJ. Interactions and activities of factors involved in the late stages of human 18S rRNA maturation. RNA Biol 2019; :1–15.

24. Wyler E, Zimmermann M, Widmann B, Gstaiger M, Pfannstiel J, Kutay U, Zemp I. Tandem affinity purification combined with inducible shRNA expression as a tool to study the maturation of macromolecular assemblies. Rna [Internet] 2011; 17:189–200. Available from: http://www.ncbi.nlm.nih.gov/entrez/query.fcgi?cmd=Retrieve&db=PubMed&dopt=Citation&list_uids=21097556

25. Sloan KE, Knox AA, Wells GR, Schneider C, Watkins NJ. Interactions and activities of factors involved in the late stages of human 18S rRNA maturation. RNA Biol 2019; 16:196–210.

26. Ferreira-Cerca S, Sagar V, Schafer T, Diop M, Wesseling AM, Lu H, Chai E, Hurt E, LaRonde-LeBlanc N. ATPase-dependent role of the atypical kinase Rio2 on the evolving pre-40S ribosomal subunit. Nat Struct Mol Biol [Internet] 2012; 19:1316–23. Available from: http://www.ncbi.nlm.nih.gov/pubmed/23104056

27. Knüppel R, Christensen RH, Gray FC, Esser D, Strauß D, Medenbach J, Siebers B, Macneill SA, Laronde N, Ferreira-Cerca S. Insights into the evolutionary conserved regulation of Rio ATPase activity. Nucleic Acids Res 2018; 46:1441–56.

28. Heuer A, Thomson E, Schmidt C, Berninghausen O, Becker T, Hurt E, Beckmann R. Cryo-EM structure of a late pre-40S ribosomal subunit from Saccharomyces cerevisiae. Elife [Internet] 2017; 6. Available from: https://www.ncbi.nlm.nih.gov/pubmed/29155690

29. Scaiola A, Peña C, Weisser M, Böhringer D, Leibundgut M, Klingauf-Nerurkar P, Gerhardy S, Panse VG, Ban N. Structure of a eukaryotic cytoplasmic pre-40S ribosomal subunit. EMBO J 2018; 37:e98499.

30. LaRonde-LeBlanc N, Wlodawer A. Crystal structure of A. fulgidus Rio2 defines a new family of serine protein kinases. Structure [Internet] 2004; 12:1585–94. Available from: http://www.ncbi.nlm.nih.gov/pubmed/15341724

31. Fung HY, Fu SC, Brautigam CA, Chook YM. Structural determinants of nuclear export signal orientation in binding to exportin CRM1. Elife [Internet] 2015; 4. Available from: http://www.ncbi.nlm.nih.gov/pubmed/26349033

32. Fischer U, Schauble N, Schutz S, Altvater M, Chang Y, Faza MB, Panse VG. A non-canonical mechanism for Crm1-export cargo complex assembly. Elife [Internet] 2015; 4. Available from: http://www.ncbi.nlm.nih.gov/pubmed/25895666

33. Wang J, Varin T, Vieth M, Elkins JM. Crystal structure of human RIOK2 bound to a specific inhibitor. Open Biol 2019; 9.

34. Ashkenazy H, Abadi S, Martz E, Chay O, Mayrose I, Pupko T, Ben-Tal N. ConSurf 2016: an improved methodology to estimate and visualize evolutionary conservation in macromolecules. Nucleic Acids Res [Internet] 2016; 44:W344–50. Available from: https://www.ncbi.nlm.nih.gov/pubmed/27166375

35. Jura N, Zhang X, Endres NF, Seeliger MA, Schindler T, Kuriyan J. Catalytic control in the EGF receptor and its connection to general kinase regulatory mechanisms. Mol Cell [Internet] 2011; 42:9–22. Available from: http://www.ncbi.nlm.nih.gov/pubmed/21474065

36. Lavoie H, Li JJ, Thevakumaran N, Therrien M, Sicheri F. Dimerization-induced allostery in protein kinase regulation. Trends Biochem Sci [Internet] 2014; 39:475–86. Available from: https://www.ncbi.nlm.nih.gov/pubmed/25220378

37. Cai Z, Chehab NH, Pavletich NP. Structure and activation mechanism of the CHK2 DNA damage checkpoint kinase. Mol Cell [Internet] 2009; 35:818–29. Available from: http://www.ncbi.nlm.nih.gov/pubmed/19782031

38. Wan PT, Garnett MJ, Roe SM, Lee S, Niculescu-Duvaz D, Good VM, Jones CM, Marshall CJ, Springer CJ, Barford D, et al. Mechanism of activation of the RAF-ERK signaling pathway by oncogenic mutations of B-RAF. Cell [Internet] 2004; 116:855–67. Available from: https://www.ncbi.nlm.nih.gov/pubmed/15035987

39. Flayhan A, Berge C, Bailo N, Doublet P, Bayliss R, Terradot L. The structure of Legionella pneumophila LegK4 type four secretion system (T4SS) effector reveals a novel dimeric eukaryotic-like kinase. Sci Rep [Internet] 2015; 5:14602. Available from: https://www.ncbi.nlm.nih.gov/pubmed/26419332

40. Larburu N, Montellese C, O’Donohue MF, Kutay U, Gleizes PE, Plisson-Chastang C. Structure of a human pre-40S particle points to a role for RACK1 in the final steps of 18S rRNA processing. Nucleic Acids Res [Internet] 2016; 44:8465–78. Available from: http://www.ncbi.nlm.nih.gov/pubmed/27530427

41. Ameismeier M, Cheng J, Berninghausen O, Beckmann R. Visualizing late states of human 40S ribosomal subunit maturation. Nature 2018; 558:249–53.

42. Sloan KE, Gleizes PE, Bohnsack MT. Nucleocytoplasmic Transport of RNAs and RNA-Protein Complexes. J Mol Biol [Internet] 2016; 428:2040–59. Available from: http://www.ncbi.nlm.nih.gov/pubmed/26434509

43. Mitterer V, Gantenbein N, Birner-Gruenberger R, Murat G, Bergler H, Kressler D, Pertschy B. Nuclear import of dimerized ribosomal protein Rps3 in complex with its chaperone Yar1. Sci Rep [Internet] 2016; 6:36714. Available from: https://www.ncbi.nlm.nih.gov/pubmed/27819319

44. Christie M, Chang CW, Rona G, Smith KM, Stewart AG, Takeda AA, Fontes MR, Stewart M, Vertessy BG, Forwood JK, et al. Structural Biology and Regulation of Protein Import into the Nucleus. J Mol Biol [Internet] 2016; 428:2060–90. Available from: http://www.ncbi.nlm.nih.gov/pubmed/26523678

45. Huber FM, Hoelz A. Molecular basis for protection of ribosomal protein L4 from cellular degradation. Nat Commun [Internet] 2017; 8:14354. Available from: https://www.ncbi.nlm.nih.gov/pubmed/28148929

46. Itzhak DN, Tyanova S, Cox J, Borner GH. Global, quantitative and dynamic mapping of protein subcellular localization. Elife 2016; 5.

47. Mitterer V, Murat G, Rety S, Blaud M, Delbos L, Stanborough T, Bergler H, Leulliot N, Kressler D, Pertschy B. Sequential domain assembly of ribosomal protein S3 drives 40S subunit maturation. Nat Commun [Internet] 2016; 7:10336. Available from: https://www.ncbi.nlm.nih.gov/pubmed/26831757

48. Kiburu IN, LaRonde-LeBlanc N. Interaction of Rio1 kinase with toyocamycin reveals a conformational switch that controls oligomeric state and catalytic activity. PLoS One [Internet] 2012; 7:e37371. Available from: http://www.ncbi.nlm.nih.gov/entrez/query.fcgi?cmd=Retrieve&db=PubMed&dopt=Citation&list_uids=22629386

49. Romier C, Ben Jelloul M, Albeck S, Buchwald G, Busso D, Celie PH, Christodoulou E, De Marco V, van Gerwen S, Knipscheer P, et al. Co-expression of protein complexes in prokaryotic and eukaryotic hosts: experimental procedures, database tracking and case studies. Acta Crystallogr D Biol Crystallogr [Internet] 2006; 62:1232–42. Available from: http://www.ncbi.nlm.nih.gov/entrez/query.fcgi?cmd=Retrieve&db=PubMed&dopt=Citation&list_uids=17001100

50. Schuck P. Size-distribution analysis of macromolecules by sedimentation velocity ultracentrifugation and lamm equation modeling. Biophys J [Internet] 2000; 78:1606–19. Available from: https://www.ncbi.nlm.nih.gov/pubmed/10692345

51. Vistica J, Dam J, Balbo A, Yikilmaz E, Mariuzza RA, Rouault TA, Schuck P. Sedimentation equilibrium analysis of protein interactions with global implicit mass conservation constraints and systematic noise decomposition. Anal Biochem [Internet] 2004; 326:234–56. Available from: https://www.ncbi.nlm.nih.gov/pubmed/15003564

52. Brautigam CA. Calculations and Publication-Quality Illustrations for Analytical Ultracentrifugation Data. Methods Enzym [Internet] 2015; 562:109–33. Available from: https://www.ncbi.nlm.nih.gov/pubmed/26412649

53. Kabsch W. Integration, scaling, space-group assignment and post-refinement. Acta Crystallogr D Biol Crystallogr [Internet] 2010; 66:133–44. Available from: http://www.ncbi.nlm.nih.gov/pubmed/20124693

54. Sheldrick GM. A short history of SHELX. Acta Crystallogr A [Internet] 2008; 64:112–22. Available from: http://www.ncbi.nlm.nih.gov/entrez/query.fcgi?cmd=Retrieve&db=PubMed&dopt=Citation&list_uids=18156677

55. McCoy AJ, Grosse-Kunstleve RW, Adams PD, Winn MD, Storoni LC, Read RJ. Phaser crystallographic software. J Appl Crystallogr [Internet] 2007; 40:658–74. Available from: http://www.ncbi.nlm.nih.gov/pubmed/19461840

56. Cowtan K. The Buccaneer software for automated model building. 1. Tracing protein chains. Acta Crystallogr D Biol Crystallogr [Internet] 2006; 62:1002–11. Available from: http://www.ncbi.nlm.nih.gov/pubmed/16929101

57. Bricogne G, Blanc E, Brandl M, Flensburg C, Keller P, Paciorek W, Roversi P, Smart OS, Vonrhein C, Womack T. BUSTER, version 2.8.0. Cambridge, United Kingdom Glob Phasing Ltd 2009;

58. DeLano WL. The PyMOL Molecular Graphics System, Version 1.8. Schrödinger LLC [Internet] 2014; :http://www.pymol.org. Available from: http://www.pymol.org

59. Emsley P, Lohkamp B, Scott WG, Cowtan K. Features and development of Coot. Acta Crystallogr D Biol Crystallogr [Internet] 2010; 66:486–501. Available from: http://www.ncbi.nlm.nih.gov/pubmed/20383002

